# The CC′ loop of IgV domain containing immune checkpoint receptors: From a bystander to an active determinant of receptor:ligand binding

**DOI:** 10.1101/656462

**Authors:** Shankar V. Kundapura, Udupi A. Ramagopal

## Abstract

Antibodies targeting negative regulators of immune checkpoints have shown unprecedented and durable response against variety of malignancies. While the concept of blocking the negative regulators of immune checkpoints using mAbs appears to be an outstanding approach, their limited effect and several drawbacks such as resistance, poor solid tumor penetration and so on, calls for the rational design of next generation of therapeutics. Soluble isoforms of negative regulators of immune checkpoints are expressed naturally and are shown to regulate the immune response, suggesting the soluble version of these molecules and affinity-modified versions of these self-molecules could be effective lead molecules for immunotherapy. To get a better insight on hotspot regions for modification, we have analysed structures of available immune receptor:ligand complexes containing IgV domains. Interestingly, this analysis reveals that the CC′ loop of IgV domain, a loop which is distinct from CDRs which are generally utilized by antibodies to recognize antigens, plays a pivotal role in affinity modulation. Here, we present several examples of cognate partner specific conformational variation observed in CC′ loop of several checkpoint receptor:ligand complexes. In addition, *in silico* swapping of CC′ loop targeting TIGIT:Nectin-2/PVR pathway corroborated well with biophysically determined affinity values for these complexes. Thus, CC′ loop appears to be a hotspot for affinity modification without affecting the specificity to their cognate receptors, an important requirement to avoid unintended interaction of these modified molecules with undesired targets.

## Introduction

The concept of blocking the negative regulators of the immune system (checkpoint blockade) to control/eradicate cancer and its durable response over the plethora of cancer types, has revolutionised the way cancer is treated today. Here, the reactivation of the immune system to fight cancer lies at the core of its philosophy. The involvement of co-stimulatory or inhibitory immune checkpoint receptors (ICRs) (Jenkins et al., 1987; Lafferty and Cunningham, 1975; Schwartz et al., 1989), along with primary MHC-2 (Major Histocompatibility Complex-2) and TCR (T-cell receptor) signalling, is paramount for the activation or deactivation of T-cells. These immune checkpoint pathways like CTLA-4/CD28:B7-1/B7-2, CD226/TIGIT:Nectin-2/PVR and PD-1:PD-L1/PD-L2, hold considerable sway over the functional fate of the T-cells, B-cells, NK cells and so on., (Pardoll, 2012). Interestingly, tumour cells have been known to exploit some of these pathways by over-expressing the ligands of the inhibitory ICRs to escape immunosurviellance (Wu et al., 2014). The blockade of signalling of these negative pathways through antibodies resulted in reinvigoration of exhausted T-cells (Freeman et al., 2006), which eventually led to regression of tumour size (Hirano et al., 2005; Iwai et al., 2002; Pilon-Thomas et al., 2010).

Extraordinary amount of research on this conceptually novel approach and consequent preclinical and clinical studies have resulted in approval of effective monoclonal antibodies (mAbs) against inhibitory ICRs, as cancers therapeutics (Goodin, 2015; USFDA, 2016a, 2017b, 2017c, 2017a), which was pioneered by the first-in-class mAb called Yervoy (Hoos et al., 2010; Wolchok et al., 2013). Immune checkpoint blockade via monoclonal antibodies became so influential in treating cancer that the Nobel prize for Physiology/Medicine 2018 was awarded to James P. Allison and Tasuku Honjo ‘*for their discovery of cancer therapy by inhibition of negative immune regulation*’. As the popularity of mAbs in cancer therapy grew, so did the reports of their drawbacks namely, high cost (Staton Tracy, 2014), poor solid tumor infiltration of these mAbs (Maute et al., 2015), moderate to severe adverse effects (Hamid et al., 2013) and the observation that the mAb based therapy is responsive only in minority of patients (Ferris et al., 2010). Further, recent findings of resistance to mAbs mediated immunotherapy is worrisome (Jenkins et al., 2018). The adverse effects observed from the clinical trials of TGN1412, an antiCD28 mAb (Attarwala, 2010; Suntharalingam et al., 2006) highlight the unpredictable consequences of such antibodies. Moreover, these antibodies have a long half-life and are administered in high dose. Hence, in the event of adverse effect of these antibodies, it is difficult to quickly withdraw them from the body (Barakat, 2014). Further, the gaps in understanding of the underlying biological mechanisms is another major setback for its logical application. For example, the observation that, distinct cellular mechanisms play a role in anti-CTLA-4 and anti-PD1 checkpoint blockade recommends further investigation on fundamental mechanisms (Wei et al., 2017). While the concept is undoubtedly a breakthrough, the above limitations suggests that we are yet to experience the full potential of immune checkpoint blockade which calls for the urgent exploration of novel modalities that can target the same pathways. There has been an encouraging shift from the use of mAbs to small molecules (Zak et al., 2016), DNA aptamers (Lai et al., 2016), peptides (Chang et al., 2015) and modified ICRs (Lázár-Molnár et al., 2017; Maute et al., 2015; Shrestha et al., 2019), some of which are in the preliminary stages of investigation and have immense potential to be on the shelf of a pharmacy in the near future.

Use of self-proteins for therapeutic purposes started with insulin and today, several such proteins are being used for various clinical applications (Leader et al., 2008). Further, the soluble versions of immune checkpoint receptors (ICRs) as Fc conjugates is a proven non-mAb based approach, which is already in clinical practice. To name a few, Orencia (Abatacept-soluble CTLA-4-IgG1Fc), Nulojix (Belatacept, a more efficacious version of Orencia) and Etanercept (commercial name Enbrel-soluble TNF receptor-IgG1Fc) are approved for treatment of moderate to severe rheumatoid arthritis (USFDA, 2016b). The success of Etanercept in treating the autoimmune disorder was widely acknowledged, so much so that it became the 3^rd^ largest selling drug in 2015 (PharmaCompass, 2016). In fact, soluble isoforms of ICRs are secreted by the immune cells to regulate the immune response (Frigola et al., 2011; Ward et al., 2013). This suggests that these isoforms which are naturally present in our body, could be rationally modified to engineer lead molecules which function similarly to mAbs. There have been interesting reports of modified ICRs which have increased binding affinity and penetrance in solid tumours when compared to mAbs. For instance, a high affinity consensus (HAC) Programmed cell death protein-1 (PD-1) mutant displayed 35,000-fold higher binding affinity to one of its ligands, PD-L1 (Programmed cell death protein-1 ligand 1) than the wild type PD-1 (Maute et al., 2015). The HAC PD-1 is reported to have better penetrance into solid tumours than the anti-PD1 mAb and displayed its potential to be a diagnostic tool for PD-L1 expressing tumours. However, this 10-point mutant was obtained from a tedious directed evolution method, which had lost its binding affinity to its other ligand PD-L2 (Programmed cell death protein-1 ligand 2). Contrary to this example, a structure based rational design of single point PD-1 mutant (A132L) displayed a 45-fold and 30-fold increased binding affinity to both its ligands, PD-L1 and PD-L2 respectively (Lázár-Molnár et al., 2017). Hence, it appears that the structure-based rational design provides an opportunity to create lead molecules with tuneable affinity without affecting the specificity to their cognate partner(s). However, for such an endeavour, a sound structural knowledge of the immune receptor:ligand complexes, is paramount.

In most of the ICRs, the membrane distal ectodomain involved in immune regulation are derived from a similar scaffold having IgV (Immunoglobulin V) domain fold. For example, in co-inhibitory molecules such as PD1, PD-L1, PD-L2, CTLA-4, CD28, B7-1, B7-2, TIM3, LAG3, TIGIT, PVR and so on, the membrane distal interacting domains are carved from an IgV domain. The IgV domain belongs to a functionally variant large superfamily called the immunoglobulin superfamily (IgSF). Nature has utilized this Ig fold to give rise to proteins with diverse functions. This makes IgSF one of the largest family of proteins, with about 2% genes of the human genome belonging to IgSF (Beck, 2003). The IgSF is broadly classified into four structural sets namely, IgC1, IgC2, IgV and IgI (Bateman et al., 1996; Harpaz and Chothia, 1994). They are differentiated based on the number of β-strands and their association and the linear residue distance between the conserved cysteines which form the canonical disulphide linkage. The Ig fold is a twisted β sandwich of roughly 100-130 amino acids, comprising of two β sheets made up of seven β-strands named A to G where, A, B and E strands form one β sheet and G, F and C strands form the other β sheet (Figure 1A-C). A canonical disulphide linkage observed between B and F strands, connects the two sheets. In almost all the cases including antibodies, IgV is the functional domain and other domains like IgC1 and IgC2 are used as the structural support. There are two additional β-strands namely, C′ and C″ present in the IgV domain (Figure 1B), which makes the front face wider, comprising of A’, G, F, C, C′ and C″ strands of an IgV domain (Figure 1D) (Natarajan et al., 2015). This domain has been found in many protein families like ICRs, antibodies, nanobodies, cell adhesion molecules and cell surface receptors (Teichmann and Chothia, 2000).

**Figure 1:**
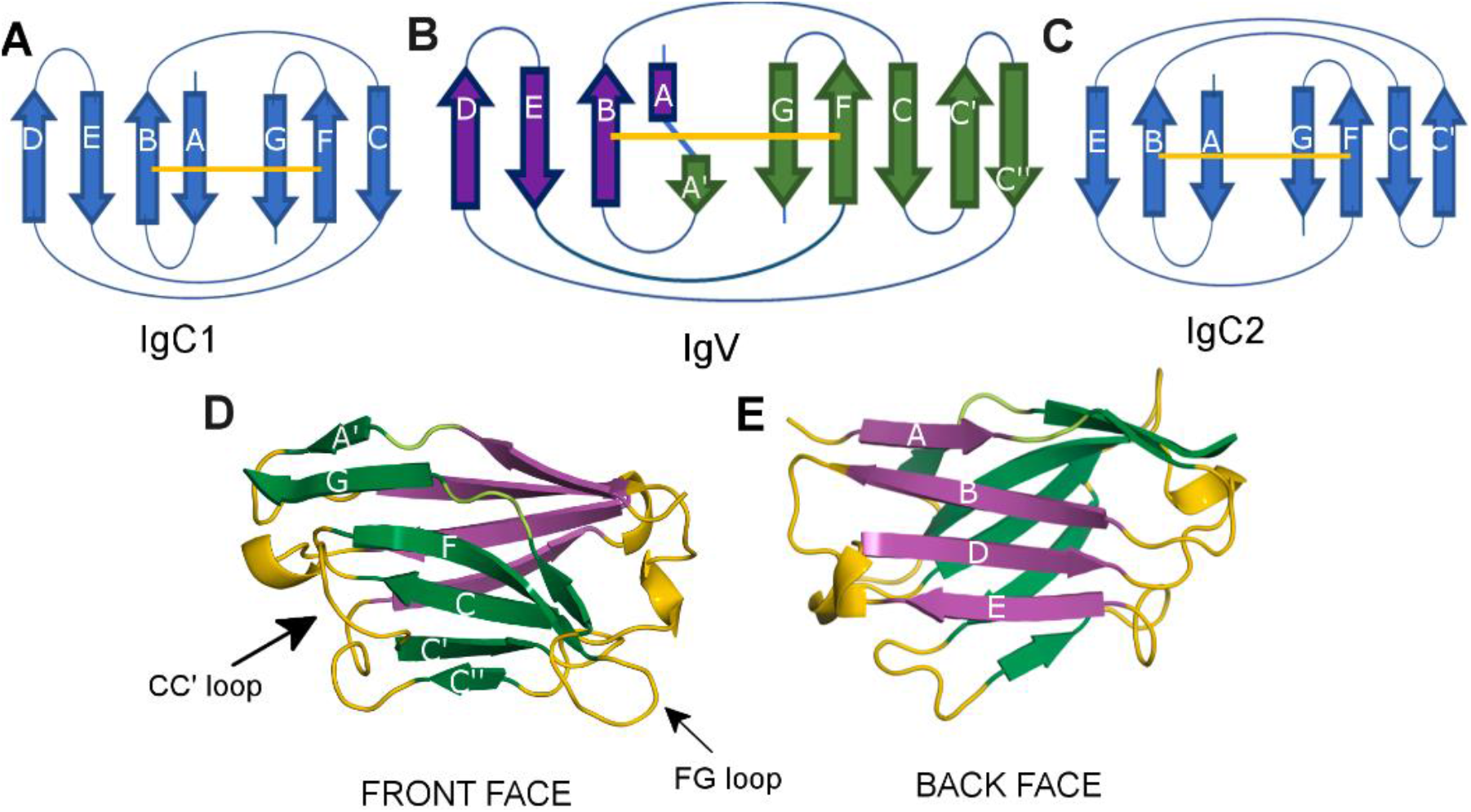
Organisation of IgV domain: Panels A, B and C illustrate the secondary structure organisation of IgC1 IgV and IgC2 domains respectively. The yellow line represents the disulphide linkage from strand B to strand F in the panels A, B and C. Panel D denotes the front face (Green ribbon) and panel E illustrates the back face (Purple ribbon) of IgV domain. Note that the front face is flanked by the FG and CC′ loops and are located at the opposite ends of the front face (Panel D).

This study aims at understanding the hotspot regions of receptor:ligand interaction to derive important insights that can be incorporated in the design of therapeutic ICRs to achieve optimal binding affinity without affecting specificity. Our goal is not directed towards the design of molecules with too many modification and exorbitant affinity to their cognate partners, as there is chance of undesirable cross-interactions with other IgV domain proteins. Instead, optimal increase in binding affinity towards the targeted ligand without sacrificing the specificity is desirable (Almo and Guha, 2014). A small, yet significant increase in binding affinity is shown to increase the biological potency of the molecule by many folds (Larsen et al., 2005). Towards this goal, we have analysed the available structure of immune receptors-ligand complexes containing IgV domain(s). It was interesting to observe that CC′ loop, which is not one of the key loops (CDRs) in IgV domain of antibodies, appears to be an active determinant of receptor:ligand interaction in the case of ICRs. To our surprise, the importance of the CC′ loop in ligand binding has been sparsely highlighted when compared to FG loop and the front face. To further authenticate our observation, TIGIT:Nectin-2/PVR pathway was considered to elucidate the importance of CC′ loop as TIGIT binds to itself and its partners in the same orientation and yet has differing binding affinities (Samanta et al., 2017), making this pathway attractive to computationally investigate the effect of modification of the CC′ loop on binding affinity. The CC′ loop of TIGIT was spliced with the CC′ loops of Nectin-2 and PVR, to investigate its effect on the binding affinity with PVR, Nectin-2 and TIGIT (homodimer).

Here, by employing structural bioinformatics tools, we found significant number of IgV domain containing ICRs which use their CC′ loops to make indispensable interactions with their cognate ligands. Overall, this study attempts to shed light on the importance of CC′ loop in few of the important IgV/Ig like domain containing ICR pathways and provide insights on effective rational modification of self-molecules for immunotherapy.

## Results and Discussion

Literature and structural search led to the identification of 18 ICR complexes which contained at least one IgV domain. Of the 18 ICR complexes, 33 IgV domains, 2 small molecules (phosphatidylserine and siallylactose) and 1 TNF superfamily member (HVEM of BTLA-HVEM complex) were present. Among them, 33 IgV domains were considered for further analysis on their mode of interaction with their cognate partners. In this analysis, the parameters used to ascertain the contribution of CC′ loop and FG loop to the interaction-interface are buried surface area (BSA) and inter-protein hydrophilic interactions. Further, post-interaction movement of CC′ loop and FG loop (C_α_ atom movement >3Å) upon binding, was taken into consideration.

### The association of IgV domains in antibodies and ICR complexes

In antibodies, each Fab (fragment of antigen binding) is a heterodimeric association of two IgV domains (termed as V_H_ and V_L_) followed by association of IgC domains of light and heavy chains. As shown in the figure 2A, the two IgV domains associate through their hydrophobic front face, where the CC′ loops (Figure 2A; shown in blue) from the V_H_ and V_L_ domains interact with each other and are a part of extended hydrophobic core. Further, the CC′ loops (Figure 2A, blue), are distal to the bound antigen; whereas the FG loops [or CDR3 of V_H_ (orange) and V_L_ (red)] located at the other end of the front face (Figure 2A) make key interaction with the antigen (Ramagopal et al., 2017). This mode of interaction of V_H_ and V_L_ domains of a Fab brings their respective variable loops (CDR1, CDR2, CDR3) in proximity to each other. These closely placed CDR loops and their variation endows ability to recognize diverse antigens with high specificity (Figure 2A) (Abhinandan and Martin, 2008; Chothia and Lesk, 1987; Chothia et al., 1998; Kabat et al., 1977).

**Figure 2:**
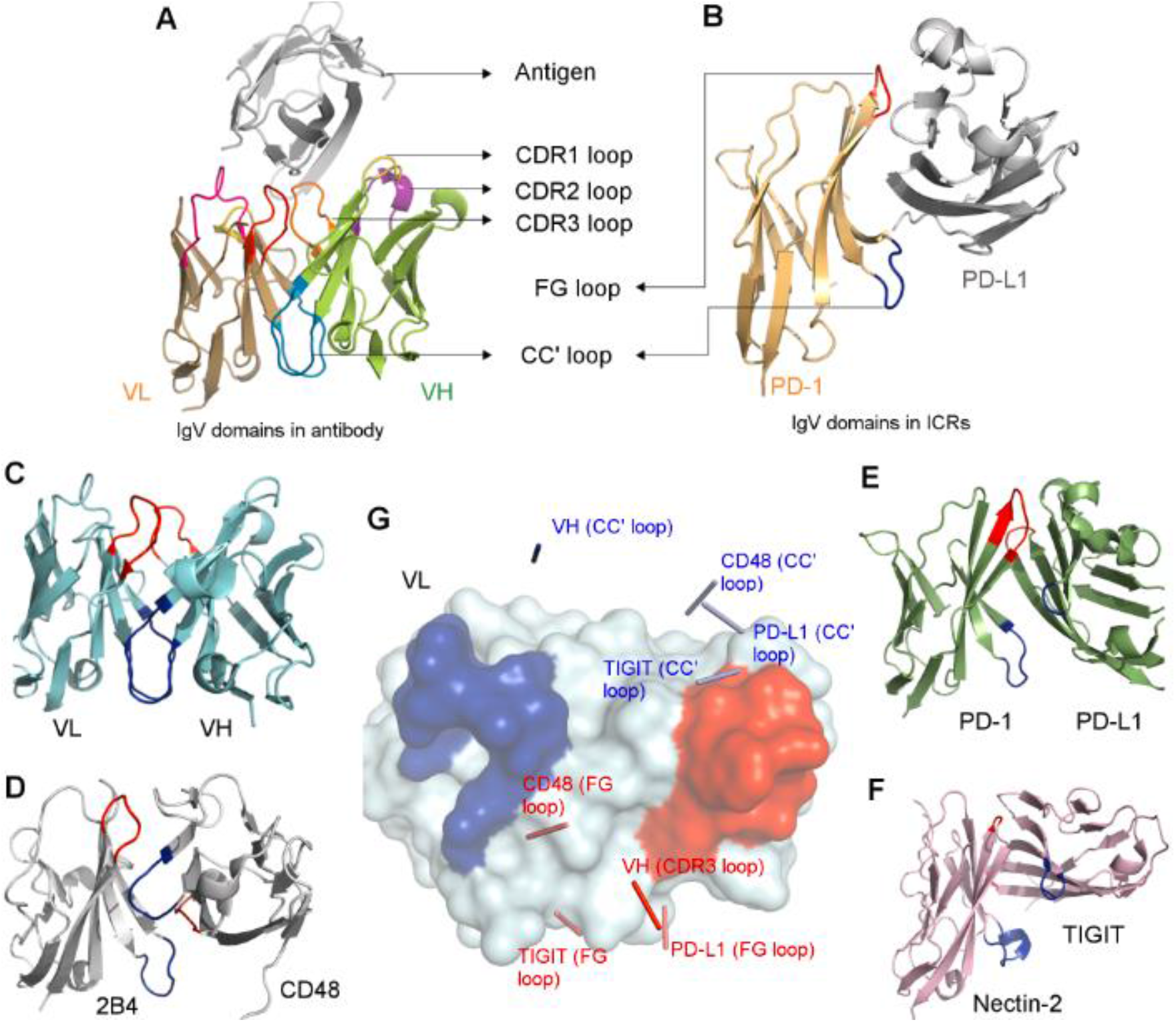
Association of IgV domains in antibodies and ICR complexes: Panel A illustrates V_H_ and V_L_ domains of an antibody presented in cartoon representation (teal), where CC′ loop is coloured dark blue (in all panels of Figure 2) and FG loop, red (in all panels of Figure 2). Panel B illustrates the 2B4:CD48 complex (grey) where positions of CC′ loops (dark blue) and FG loop (red) are marked. The same is illustrated in PD-1:PD-L1 (forest green) shown in panel C and TIGIT:Nectin-2 (deep salmon) shown in panel D. The V_L_ domain (surface, teal, panel A) of the antibody was superposed with PD-1 of PD-1:PD-L1 complex, Nectin-2 of the TIGIT:Nectin-2 complex, 2B4 of the 2B4:CD48 complex. The position of CC′ loops and FG loops of V_H_ domain, PD-L1, TIGIT and CD48 are marked by ribbon representation, shown as thin rods connecting two central residues of the respective loops (Panel E). The arrangement of the CDR loops and CC′ loops of the V_H_ and V_L_ domains of antibodies is shown in panel F, where only CDR loops interact with the antigen. Panel G illustrates the positions of CC′ loop and FG loop of the IgV domain of ICR (PD-1:PD-L1) where the FG loop, front face and CC′ loop make key interactions with its cognate ligand.

However, in significant number of ICRs and other receptors, these IgV domains are mostly present as a single chain, where it is the only one N-terminal IgV domain which interacts with their cognate partner. Consequently, the front face is relatively polar in IgV domains of ICRs, and predominantly utilizes the membrane distal FG loop (corresponding to CDR3 loop in antibodies), the flat front-face comprising of G, F, C, C′ and C″ (and A’ strand in most cases) strands and the membrane proximal CC′ loop for binding to their respective binding partner (Figure 2B), unlike the IgV domains of antibodies which use predominantly the CDR loops to recognize an antigen.

Although, the association of two IgV domains (V_H_ and V_L_) in antibodies and those seen in ICRs complexes are mediated by the front face and the mode of their interaction is considerably similar, our analysis suggests that several aspects of their association are quite different from each other. In antibodies, the CC′ loop from two associating IgV domains are always close to each other and are involved in the formation of heterodimeric hydrophobic core, which restricts their relative orientation of two IgV domains. Interestingly, no such unique orientation is observed in the interactions of IgV domains of different ICR complexes. In fact, almost every ICR complex has a different binding orientation where the positions of CC′ loops and FG loops are variable (Figure 2C-G). This is illustrated well in the case of m2B4-mCD48 complex (PDB ID: 2PTT) where the CC′ loop of mCD48 is found near the vicinity of the FG loop of m2B4 and vice versa (Figure 2D, 2G). In some cases, for example homodimeric NTB-A, two CC′ loops are close each other, whereas the FG loops are far away (Cao et al., 2006). These different binding orientations observed in ICRs point to the contribution of CC′ loop towards the binding affinity of the complex. In ICR complexes, the role of the front face, FG loop and the CC′ loop are much like the role of palm (front face), index-finger (FG loop) and thumb (CC′ loop) of a clapping hand. Hence, the FG loop and the CC′ loop flanking the front face act as clamps in order to hold the incoming binding partner. This analysis also suggests that the CC′ loop makes vital interactions that fine-tune the affinity between the interacting IgV domains, whereas the FG loop and the front face provide the required specificity.

Based on these observations we analysed 18 ICR complexes and it was observed that the CC′ loop of about 49% of the proteins analysed, contributed more than 10% to the total BSA of interaction interface of one IgV domain. This is a significant statistic as the average length of the CC′ loop is only 5 residues long. Hence, it is interesting to observe how a 5-residue long loop in a ~120 residue protein contributes more than 10% of the total BSA of the interaction interface in nearly half of the ICR complexes analysed. Additionally, along with the key BSA contributions, about 27% of the 33 proteins analysed, had hydrophilic interactions like hydrogen bonds and salt bridges contributed by the CC′ loop towards the interaction interface. It was also observed that about 24% of the proteins considered for this study, had more BSA contribution from CC′ loop than the FG loop towards the interaction interface (Table 1). These statistics hints at the considerable influence of CC′ loop in the interaction-interface of these complexes.

**Table 1:**
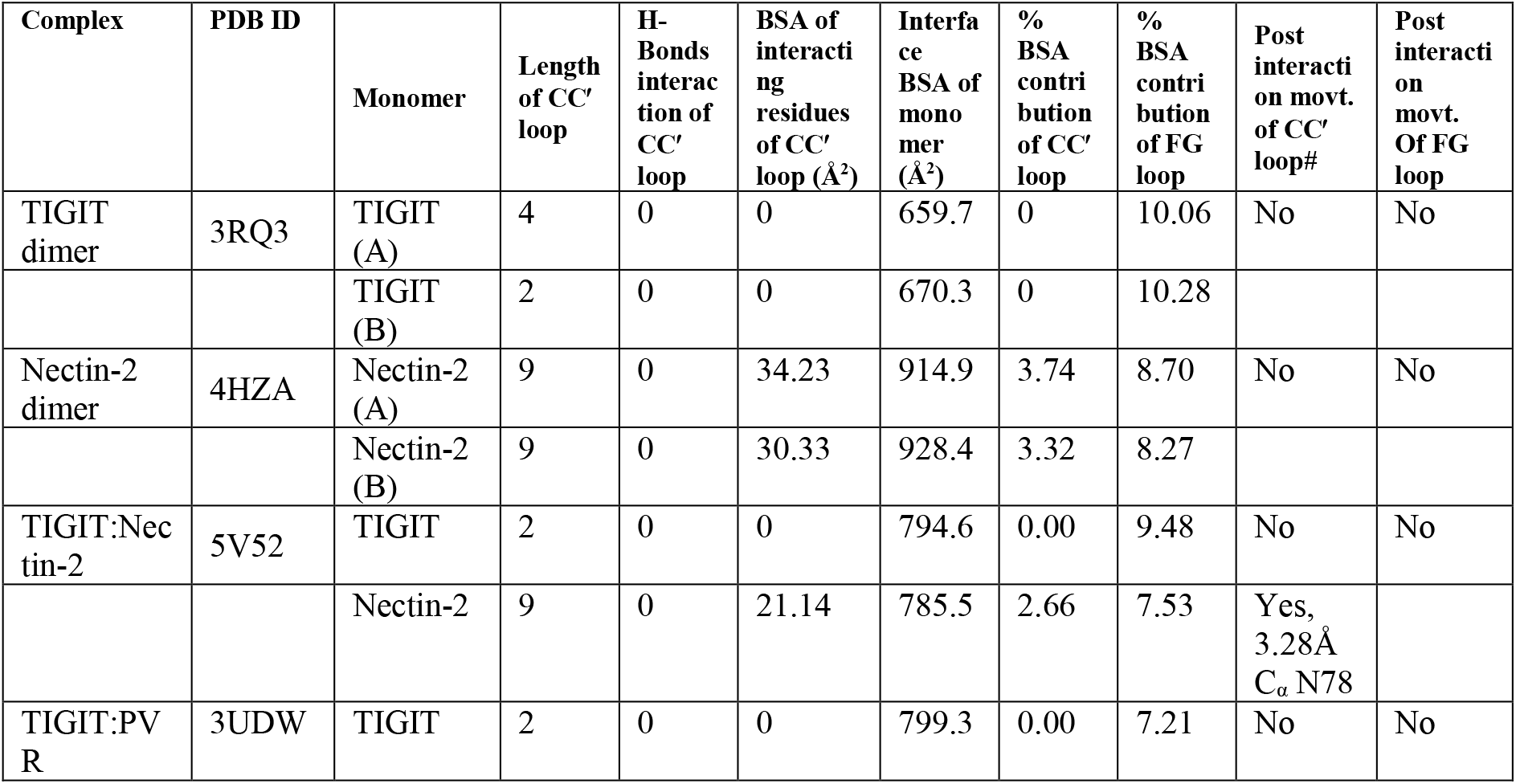
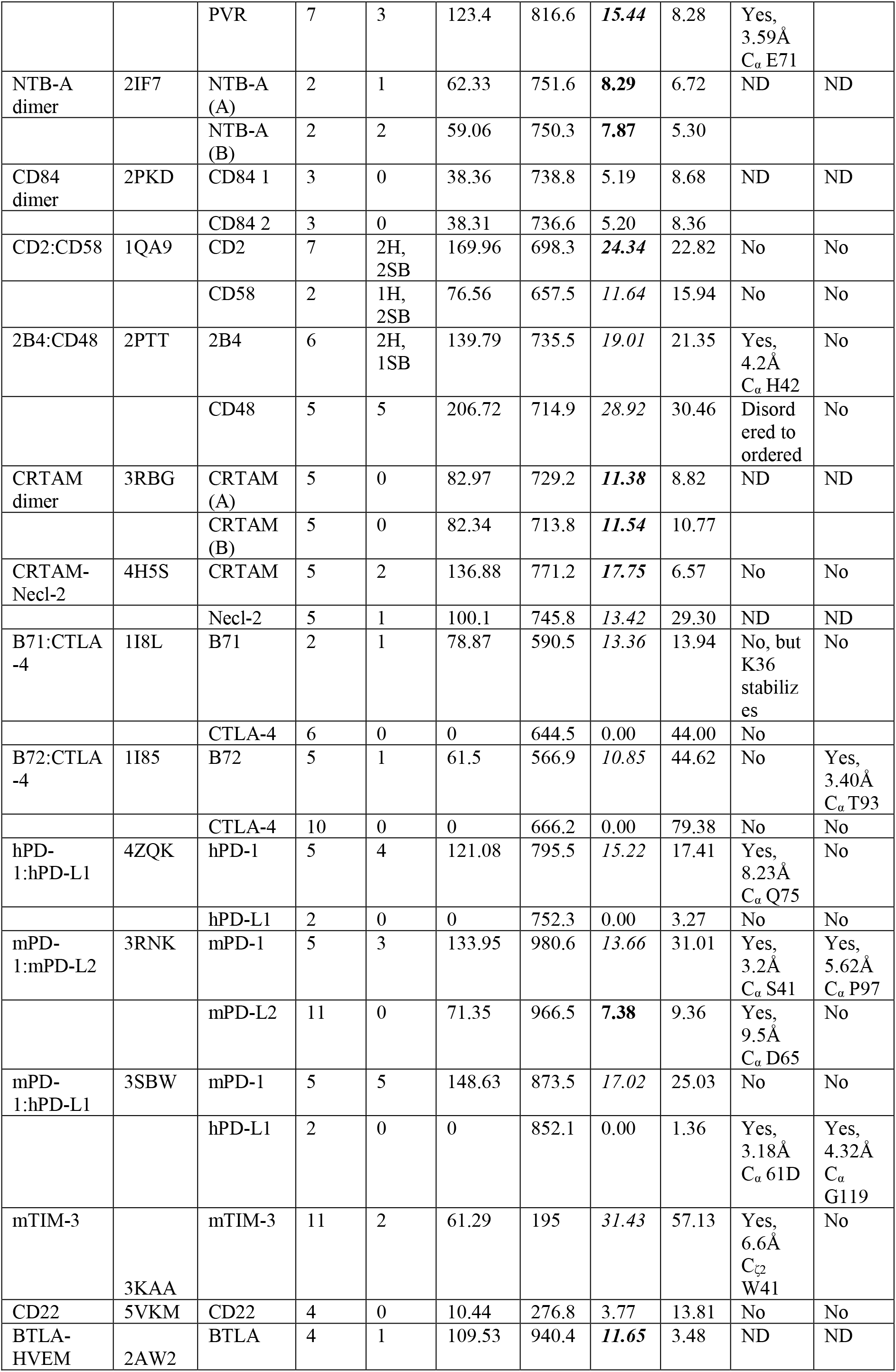
Interaction analysis of 18 IgV domain containing ICR complexes. Entries in italics indicate 10% or higher BSA contribution of the CC′ loop to the interaction-interface. Entries in BOLD denote BSA contribution of CC′ loop higher than that of FG loop. Note that the residues that are labelled as loop in PDB header file and which connects the C and C′ strands are considered as CC′ loop. The same was considered for FG loop. # post of interaction movement of any C_α_ atoms of CC′ loop and FG loop more that 3Å is considered as significant. Movement is abbreviated as Movt. And Not determinable is abbreviated as ND.

Post-interaction conformational changes or the induced fit of CC′ loop was another interesting feature that was observed. About 27% of the analysed ICRs showed deviation of more than 3Å, of the C_α_ atom of at least one of the residues in the CC′ loop before and after interaction. This value is only 9% in the case of FG loop. Further, the extent of post-interaction movement of CC′ loop is significantly larger when compared to that of the FG loop where, maximum deviation of 9.5Å and 5.6 Å have been observed for CC′ loop and FG loop respectively. This difference seems to indicate that the CC′ loop is subjected to more post-interaction movements to accommodate the incoming ligand than FG loop. It must be pointed out that alongside front face, the FG loop appears to play a pivotal role in providing the specificity of the ICR interactions. This is observed in TIGIT:TIGIT/Nectin-2/PVR pathway where a conserved ^112^TYP^114^ motif present on the FG loop of TIGIT helps in the lock-and-key interaction with its ligands Nectin-2 (Samanta et al., 2017), Polio virus receptor(PVR) and itself (Stengel et al., 2012). Similarly, the FG loop of hCTLA-4 and hCD28 with a rigid ^99^MYPPPY^104^ motif provides specificity to their cognate receptors, B7-1 and B7-2 (Schwartz et al., 2001). Hence, it could be hypothesized while FG loop is involved in conferring specificity, CC′ loop’s ability to undergo ligand induced conformational changes might be contributing to additional affinity.

As depicted in the table 1, it is evident that the CC′ loop appears to contribute significantly to ligand:receptor interactions in ICR complexes and in some cases, ligand induced conformational changes appear to be an important factor. Hence, based on the role of CC′ loop and intricacies of the post-interaction conformation changes in the CC′ loop, some interesting features were highlighted, 1) Significant post-interaction movement of CC′ loop 2) Loop to β-strand transition of CC′ loop upon interaction 3) Unusual conformation of CC′ loop leading to non-canonical IgV domain. In the following sections we provide few examples of post-interaction conformational changes and two examples of significant variation in CC′ loop leading to non-canonical conformation that carve a binding pocket for small molecules.

#### 1. Significant post-interaction movement in CC′ loop

In murine 2B4:CD48 complex, 2B4 is a member of signalling lymphocyte activation molecule (SLAM), and a receptor expressed on NK cells. SLAM family members are considered as important targets for anti-myeloma immunotherapy (Radhakrishnan et al., 2017). 2B4 positively regulates NK cell function and makes heterophilic interaction with CD48 (Velikovsky et al., 2007). This complex is involved in Interleukin-2 associated propagation and activation of NK cells which leads to NK cell mediated cytotoxicity and tumour clearance (Lee et al., 2006). The CC′ loop of the m2B4 comprising of residues ^38^EQGSHR^43^ (PDB:2PTT) (Velikovsky et al., 2007) contributes 19.01% to the total 2B4 interaction-interface BSA. This CC′ loop is also seen to contribute two hydrogen bonds and one salt bridge (Figure 3B). Furthermore, the CC′ loop of CD48 comprising of ^37^TKNQK^41^ (PDB:2PTT) contributes 28.92% to the total CD48 interaction-interface BSA, along with 5 hydrogen bonds (Figure 3C). It appears that the observation of post-interaction movement of CC′ loop of 2B4 and CD48 is vital in this context (Figure 3A). The CC′ loop of 2B4 is reported to be flexible in its apo form, among the 4 molecules in the asymmetric unit, this loop is not modelled in 2 molecules due to its disordered state. However, upon interaction, the CC′ loop of 2B4 moves to accommodate CD48 and makes key Van der Waals interactions and hydrophilic interactions, as mentioned earlier. This movement is characterized by the maximum deviation of about 4.2 Å of C_α_ atom of H42, when apo (PDB:2PTU) and bound 2B4 (PDB:2PTT) were compared (Figure 3A). The CC′ loop of apo CD48 is highly flexible and was not modelled in any of the molecules in its asymmetric unit. However, upon complexation, the CC′ loop assumes a rigid hook like conformation which is indicative of vital interactions with 2B4 (Figure 3A). This is a unique feature where CC′ loops of both 2B4 and CD48 show post-interaction movements unlike FG loops of these proteins, which contribute 28.51% (2B4) and 30.89% (CD48) to the total interaction-interface BSA. Since, the CC′ loops of 2B4 and CD48 are located on opposite sides of each other, the post-interaction movement of both the CC′ loops seem to act like clamps to increase the binding affinity of the 2B4:CD48 interaction.

**Figure 3:**
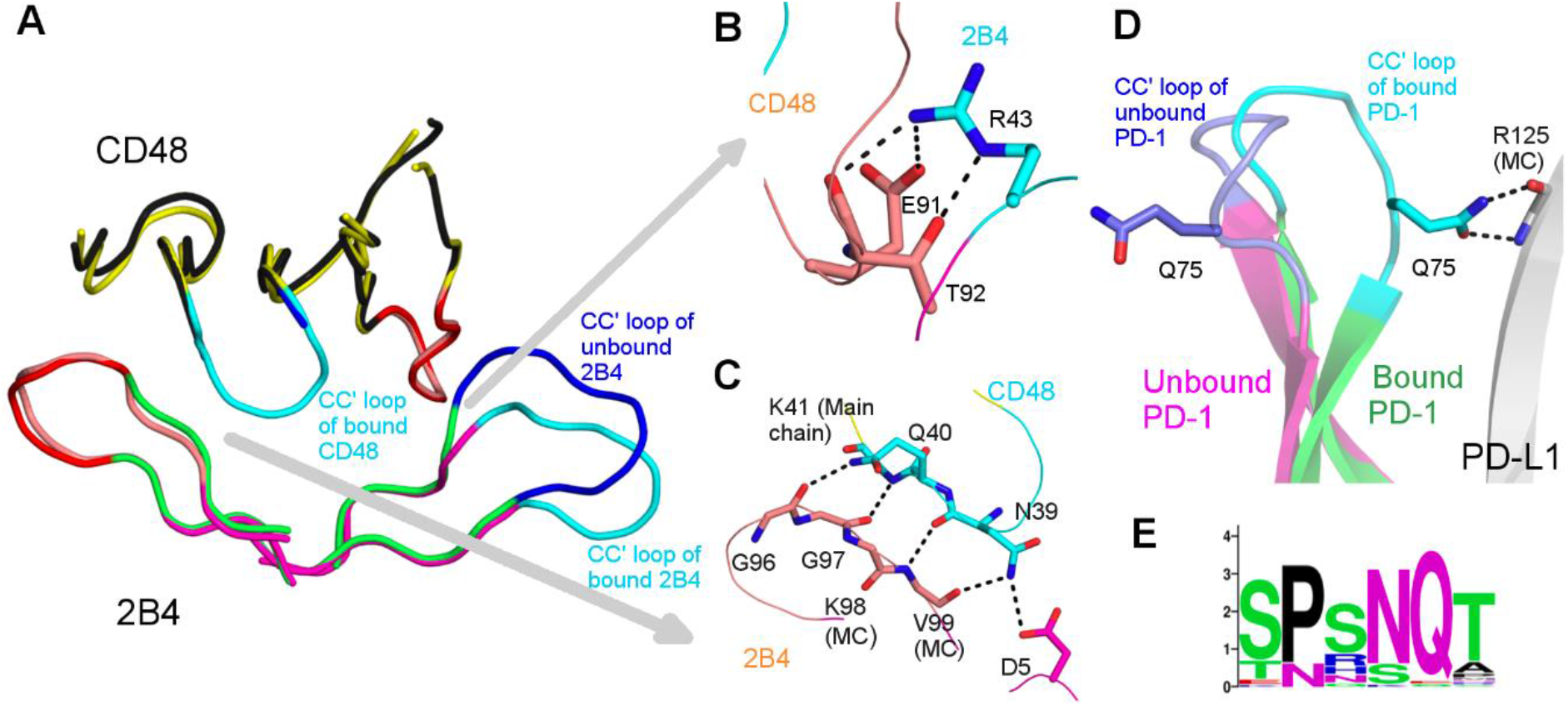
Significant post-interaction movement in CC′ loop: Panel A illustrates the superposition of 2B4 (ribbon, black):CD48 (ribbon, magenta) complex with apo structures of 2B4 (ribbon, yellow) and CD48 (ribbon, green), where CC′ loops of bound 2B4 and CD48 is shown in cyan and unbound 2B4 is shown in dark blue (only relevant part is shown for clarity). The CC′ loop of apo CD48 is not represented as it is not modelled in the PDB structure (PDB:2PTV). The FG loops of bound 2B4 and CD48 is shown in salmon and that of the apo structures is shown in red. Panel B illustrates the interactions of R43 (stick, cyan) of the CC′ loop of 2B4 with CD48 (stick, salmon). Panel C illustrates the interactions of the CC′ loop of CD48 (stick, cyan) with 2B4 (stick, salmon). Panel D illustrates the post-interaction movement of the CC′ loop (cyan) of PD-1 (cartoon, green) upon interaction with PD-L1 (cartoon, grey) when compared with the CC′ loop (dark blue) of apo PD-1 (cartoon, magenta). The sequence logo of residues of the CC′ loop of PD-1 is depicted in panel E.

A similar significant post-interaction movement of the CC′ loop is seen in human PD-1 (hPD-1) when it interacts with human PD-L1 (hPD-L1) (Zak et al., 2015). PD-1 is a well-studied inhibitory cell surface receptor (Freeman et al., 2000) inducibly expressed on T-cells, NK cells, B-cells, dendritic cells and macrophages (Agata et al., 1996; Keir et al., 2008; Yamazaki et al., 2002). Its ligands PD-L1 and PD-L2 have been observed to be over-expressed on several kinds of tumour cells to escape immunosurveillance (Ghebeh et al., 2006; Hamanishi et al., 2007; Yearley et al., 2017). The CC′ loop of hPD-1 comprising residues ^71^SPSNQ^75^ (PDB:4ZQK) contributes 15.22% to the total PD-1 interaction-interface BSA and makes four hydrogen bonds with PD-L1. Q75 of PD-1 is observed to contribute a BSA of 98.29 Å^2^ to the interaction-interface and makes two side-chain to main-chain hydrogen bonds with R125 of PD-L1 and one side-chain to side-chain hydrogen bond with D26 of PD-L1. Interestingly, a large conformational change is observed in the CC′ loop of PD-1 where a maximum deviation of 8.23 Å in C_α_ atom of Q75 is observed before and after its interaction with PD-L1 (Figure 3D). In contrast, there is no post-interaction movement of the FG loop of PD-1 which contributes 17.41% of the total PD-1 interaction-interface BSA. Likewise, no post-interaction movement is seen in the CC′ loop and FG loop of its interacting partner PD-L1. Unlike 2B4:CD48, there is only one-sided clamping of the CC′ loop of PD-1. It is also interesting to note that, of all the residues comprising the CC′ loop, only Q75 is completely conserved across most species (Figure 3E). This opens the possibility to modify other residues of CC′ loop to enhance the binding affinity, which might aid in producing lead molecule for therapeutic and diagnostic purpose.

#### 2. Loop to β-strand transition of CC′ loop upon interaction

PD-L2 is another ligand which binds to PD-1 with 3 times higher affinity than PD-L1(Cheng et al., 2013). A unique feature in mPD-L2 is observed where, the part of CC′ loop of mPD-L2 undergoes a structural transition from a loop into a β strand conformation, upon binding with its partner mPD-1 (Figure 4A). The CC′ loop of apo mPD-L2 comprises of residues ^62^VENDTSLQSERA^73^ (PDB: 3BOV) (Lázár-Molnár et al., 2008), however upon binding to mPD-1, ^67^SLQ^69^ of the CC′ loop assumes a β strand conformation (PDB:3BP5) (Lázár-Molnár et al., 2008) (Figure 4B). This could be attributed to the stabilization of the long CC′ loop of mPD-L2 after binding to mPD-1. The conformational change observed in the CC′ loop of PD-L2 is accompanied by a large movement of the CC′ loop of mPD-L2 towards mPD-1, upon binding to mPD-1, which is represented by a deviation of 9.5Å of C_α_ atom of D65 of the CC′ loop of mPD-L2. However, for such a large post-interaction movement, the contribution of the CC′ loop to the total PD-L2 interaction-interface is quite low (7.38%) and we did not observe any hydrophilic interactions. A similar feature of post interaction movements of CC′ loop and FG loop is observed in mPD-1, where a deviation of 3.2Å in the C_α_ atom of S41 and a deviation of 5.62Å in the C_α_ atom of P97, was observed in CC′ loop and FG loop respectively (PDB:3BP5). The CC′ loop and FG loop of mPD-1 contributes 13.66% and 31.01% towards the interaction-interface BSA of mPD-1. The post-interaction movements of CC′ loops of mPD-1 and mPD-L2 and FG loop of mPD-1 causes a three-way clamping which might contribute to the increased binding affinity (Figure 4C). Hence, rational and structure-based modification in any of these loops to enhance the hydrophobic or hydrophilic interactions, might possibly increase the binding affinity. Little analysis and hence, hypothesis could be given for the human counterpart, as structures of apo human PD-L2 and complex of human PD-1 and PD-L2 have not been reported thus far.

**Figure 4:**
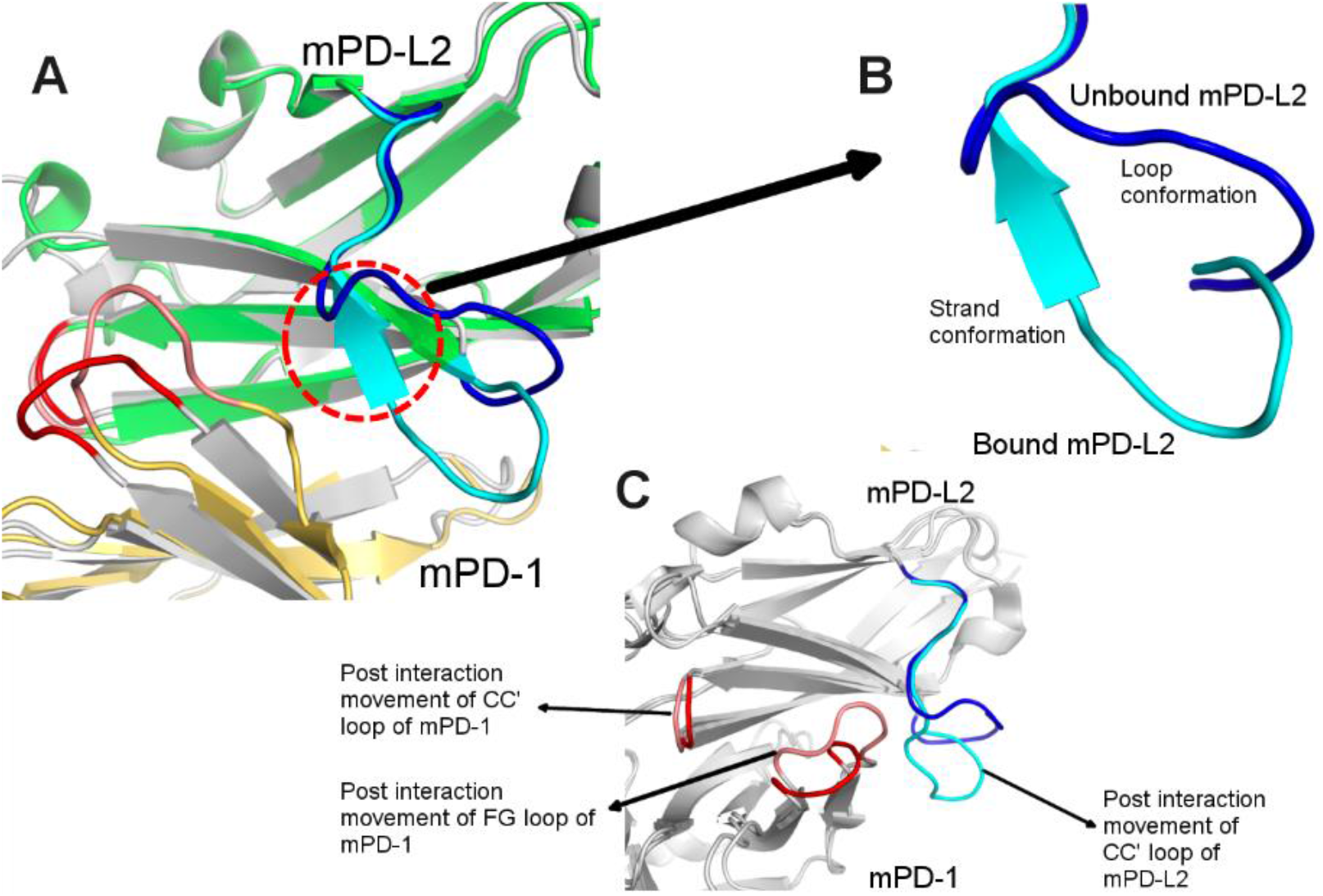
Loop to β-strand transition of CC′ loop: Panel A illustrates the superposition of mPD-1 (cartoon, gold):mPD-L2 (cartoon, green) complex with apo structures of mPD-1 (light grey) and mPD-L2 (grey), only the relevant part is shown for clarity. The CC′ loops of bound and unbound mPD-L2 is cyan and dark blue and FG loops are maintained in the same manner as figure 3. Panel B depicts the loop to β strand transition of CC′ loop of mPD-L2 upon binding to mPD-1. Panel C highlights the post-interaction movement of the CC′ loop of mPD-L2 and FG loops of mPD-1 and mPD-L2 which causes a three-way clamping.

#### 3. Unusual conformation of CC′ loop leads to non-canonical IgV domain

The examples of ICR complexes involving IgV domains described in the previous sections are protein-protein complexes. There are a few examples of immune receptors having IgV domains that recognize biologically important small molecules, suggesting the versatility of this domain to carve its front face to recognize molecules as big as proteins and small molecules of size of an amino acid. The relatively flat canonical front face of the IgV/Ig like domain does not seem to support a cleft that can accommodate binding of small molecules. Hence, for the IgV domain to acquire such a property, it appears that a non-canonical conformational change is required to accommodate small molecules (Figure 5A). This non-canonical conformation change was observed for the first time in the case of murine T-cell immunoglobulin and mucin domain containing protein-3 (mTIM-3), where unusual conformation of the CC′ loop creates a cleft for small molecule binding (Cao et al., 2007). mTIM-3 is primarily expressed on Th1 and dendritic cells (Rodriguez-Manzanet et al., 2009). They bring about apoptosis in Th1 cells via downstream inhibitory signalling, when phosphatidylserine (PtdSer) expressed on cells destined for apoptosis, bind to mTIM-3 (Zhu et al., 2005). As depicted in the figure 5A and 5C, the CC′ loop of mTIM-3 is peeled out of the IgV domain core towards FG loop of mTIM-3, forming a narrow pocket between CC′ loop and FG loop, which is known to bind PtdSer. As explained in the introduction, in a canonical IgV/ Ig domain, the tip of CC′ loop and FG loop are separated approximately by 25 Å and these two loops are located at the two ends of the IgV domain (Figure 2B, 5A). This binding pocket carved between the CC′ and FG loops is conserved in all TIM family proteins (Cao et al., 2007; Santiago et al., 2007a, 2007b). Note that such unusual peeling of C and C′ strands as well as the connecting loop away from the core would fundamentally require (1) change in the side-chain functionality of the core forming hydrophobic residues which are exposed to the solvent due to this non-canonical conformation (2) a strong interaction that can peel the CC′ loop away from its canonical position to form a pocket with FG loop. As expected, this unusual conformation of CC′ loop is stabilized by two non-canonical disulphide bonds between C52 of the C strand with C63 of the CC′ loop and C110 of F strand with C58 of the CC′ loop (PDB:3KAA) (Figure 5C) (DeKruyff et al., 2010). These non-canonical disulphide bonds pull the CC′ loop from its canonical position and away from the core towards the FG loop. Further, few of the core-forming residues of the C and C′ strands and those from the core that interact with these residues in canonical IgV domains, are changed to hydrophilic residues. This pocket is further stabilized by few hydrogen bonds and salt-bridges.

**Figure 5:**
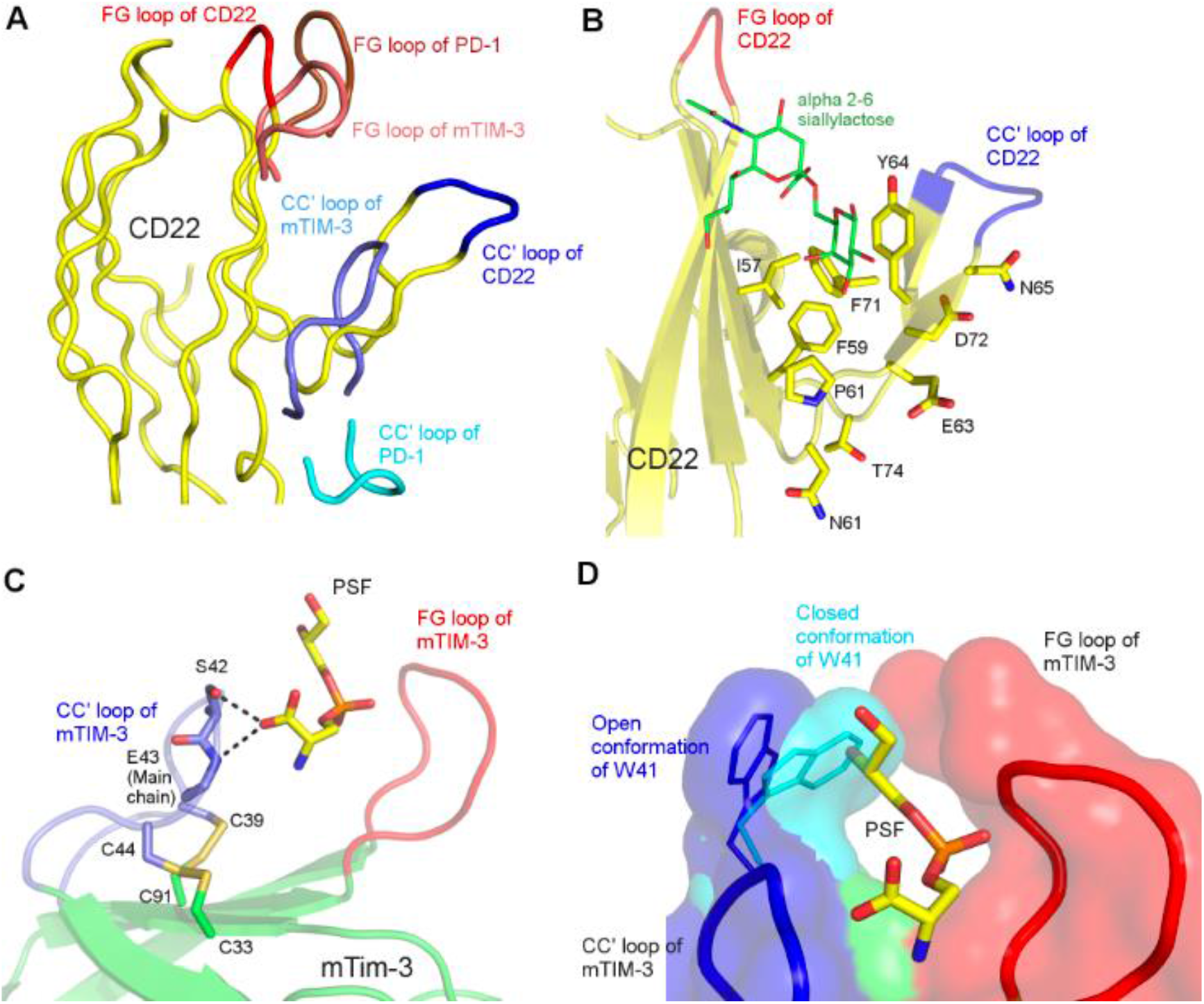
Unusual conformation of CC′ loop leads to non-canonical IgV domain: Panel A illustrates the peeling of CC′ loop (ribbon, dark blue) of CD22 (ribbon, yellow) and CC′ loop (ribbon, Slate) of mTIM-3 in comparison with the canonical CC′ loop of hPD-1 (ribbon, cyan), from the core to form a cleft between FG loop and CC′ loop. This non-canonical conformation of CC′ loops of CD22 and mTIM-3 creates a cleft between the CC′ and FG loops (CD22 (ribbon, red) and mTIM-3 (ribbon, salmon)). Panel B illustrates the pocket formed by CC′ loop (dark blue) and FG loop (red) of CD22, with bound α2-6 siallylactose (sticks, green). The residues responsible for this non-canonical CC′ loop conformation is shown in sticks (yellow). Panel C illustrates the non-canonical conformation of CC′ loop (slate) which is stabilised by 2 non-canonical disulphide linkages (shown in sticks) in mTIM3. The CC′ loop forms a narrow pocket with FG loop of mTIM-3 (red), where PtdSer (sticks, yellow) is shown to interact. Panel D illustrates the gating of the above-mentioned pocket in mTIM-3 by W41, where W41 on CC′ loop assumes a closed conformation (stick and surface, cyan) and is present close to FG loop. W41 is flung away from the FG loop to assume an open conformation (stick and surface, dark blue) to allow the entry of PtdSer.

The CC′ loop of mTIM-3 comprising of ^55^KGFCPWSQCTN^65^ (PDB:3KAA) contributes about 31.43% to the total mTIM-3 interaction-interface BSA and makes two potential hydrogen bonds with the serine moiety of PtdSer (Figure 5C). W41, present on the tip of the CC′ loop acts as a gate for this pocket, regulating the entry of PtdSer. W41 in the closed conformation connects the CC′ loop and FG loop via hydrophobic interactions (PDB:2OYP) (Cao et al., 2007). However, in the presence of PtdSer, W41 flips and rotates in such a manner that it moves away from the FG loop and allows PtdSer to enter the pocket (Figure 5D). Mutagenesis studies suggested that mutation of ^60^WSQ^62^ to homologous human sequence of VFE critically reduced the binding affinity of mTIM-3 to PtdSer (DeKruyff et al., 2010). These observations and experiments emphasize the role of this non-canonical CC′ loop in forming the pocket for an IgV domain to interact with a small molecule and plays a role of gating the entry of the small molecule into the pocket.

Another important example of CC′ loop of the IgV domain assuming a non-canonical conformation to form a small molecule binding cleft is that of CD22. CD22 is receptor present on B-cells and maintains a basal level of inhibition of B-cell activation. CD22 is a member of the sialic acid-binding immunoglobulin-like lectins family (Siglecs), found only on B-cells (Walker and Smith, 2008) and binds specifically to α2-6 siallylactose. Since, CD22 is purported to bind to α2-6 siallylactose, a non-canonical feature is observed similar to that of mTIM-3, where the C, C′ strands and CC′ loop move away from the core to form a unique pocket (PDB:5VKM) (Ereño-orbea et al., 2017). The C and C′ strands of CD22 are bent towards the N-terminal forming a β-hairpin. This β-hairpin structure along with the FG loop forms a comparatively wide but shallow pocket, where α2-6 siallylactose binds to CD22 (Figure 5B).

As mentioned earlier, the non-canonical conformation of the CC′ loop to form a pocket with FG loop could be due to the presence of hydrophobic residues like I57, F59, P62 and Y64 on the C strand and Y71 on C′ strand in CD22 which forms a hydrophobic core on the front face (Figure 5B). This hydrophobic core enables the C and C′ strand to move towards the FG loop. Furthermore, the presence of polar residues like N61, E63 and N65 on the C strand and D72 and T74 on the C′ strand, which would be facing the hydrophobic core of a canonical IgV domain, might have pushed the C and C′ strand away from the protein hydrophobic core due to steric clashes.

In a recent study that describes the structure of CD160:HEVM receptor complex (PDB:6NG3, manuscript in press), it was observed that a similar reorientation of C and C′ strands together with a single non-canonical disulphide bridge creates a pocket in the front-face of CD160, which is wider than that is observed in TIM-3 and CD22. The dimension of the HVEM receptor being slightly smaller than the IgV domains, and obviously wider than the small molecules, requires complementary surface on CD160 that would exactly fit the dimensions of the HVEM receptor. It is interesting to note that unlike TIM family members with two disulphide bridges on the front-face, how a relative wider surface is created using single disulphide bridge. Furthermore, point mutations on the CC′ loop of CD160, resulted in the reduced binding to HVEM receptor. All these results suggest that how nature uses the versatility of C and C′ strands together with CC′ loop to create ligand binding surface of different dimensions.

A question piqued our interest in these structures, why does the CC′ loop assume a non-canonical conformation to form a pocket? Why doesn’t the onus fall on FG loop to form this non-canonical conformation? Upon closer observation, it might be due to the presence of the canonical disulphide bond between the B strand and F strand of the IgV domain. This covalent bond might restrict the flexible movement of the FG loop and/or F strand. Hence, we propose that, since CC′ loop is not constrained by the presence of any covalent bonds or other rigidity conferring interactions, it is a prime candidate in the front face to assume non-canonical conformations to incorporate and provide additional affinity to its respective binding partner.

### Modification of CC′ loop might lead to increase in binding affinity: A case study of TIGIT/Nectin pathway

T-cell immunoreceptor with Ig and ITIM domain (TIGIT) is an inhibitory receptor expressed on NK cells and certain subset of T-cells. TIGIT belongs to the PVR/Nectin family (Yu et al., 2009) and is responsible for blocking the anti-tumor activity in NK cells (Vales-Gomez et al., 1998) and T-cells (Yu et al., 2009). TIGIT binds to itself, PVR and Nectin-2 in a canonical mode where the Y/FP motif present on the FG loop of all the three proteins recognize a structurally conserved complementary cleft on the front face of their partners (Deuss et al., 2017; Samanta et al., 2017; Stengel et al., 2012). Superposition of TIGIT:TIGIT, TIGIT:Nectin-2 and TIGIT:PVR structures, clearly suggest TIGIT binds to all its binding partners and to itself in the same manner due to the, so called ‘lock and key’ mode of interaction, but with differing binding affinity (Samanta et al., 2017). TIGIT binds to itself to form a very weak homodimer which could be detected at 350μM, also binds to Nectin-2 at 11.2μM and has the highest binding affinity towards PVR at 0.35μM (Samanta et al., 2017). Among a few factors for these varied binding affinities between TIGIT and its ligands, the differing lengths and side-chain functionalities of the residues of CC′ loops of TIGIT, Nectin-2 and PVR (Figure 6A) was proposed as a key factor. This similarity in the mode of association of TIGIT with itself and with Nectin-2 and PVR gives us an opportunity to computationally investigate the impact of the CC′ loop on the binding affinity. Considering these observations, TIGIT/Nectin pathway presents itself as a prime example for investigating such a hypothesis. The CC′ loop of TIGIT is the shortest with 4 polar residues (^61^QQDQ^64^) (PDB:3RQ3), Nectin-2 has the longest CC′ loop with 9 residues (^72^RPDAPANHQ^80^) (PDB:5V52) (Deuss et al., 2017) and PVR has a 7-residue long CC′ loop (^69^HGESGSM^75^) (PDB:3UDW) (Stengel et al., 2012). The CC′ loop of TIGIT makes no contribution to the interaction interface of the TIGIT homodimer, slight Van der Waals interactions are provided by the residues of CC′ loop of Nectin-2 towards the interaction interface of TIGIT:Nectin-2 heterodimer. However, a significant mix of Van der Waals and H-bond interactions are provided by the CC′ loop of PVR towards the interaction interface of TIGIT:PVR heterodimer (Table 1) (Figure 6B-D). Since, TIGIT forms a weak homodimer and the contribution from its CC′ loop is low, could the binding affinity of TIGIT with itself be increased by splicing its CC′ loop with that of the CC′ loops of PVR and Nectin-2 on one of the protomer of the TIGIT homodimer?

**Figure 6:**
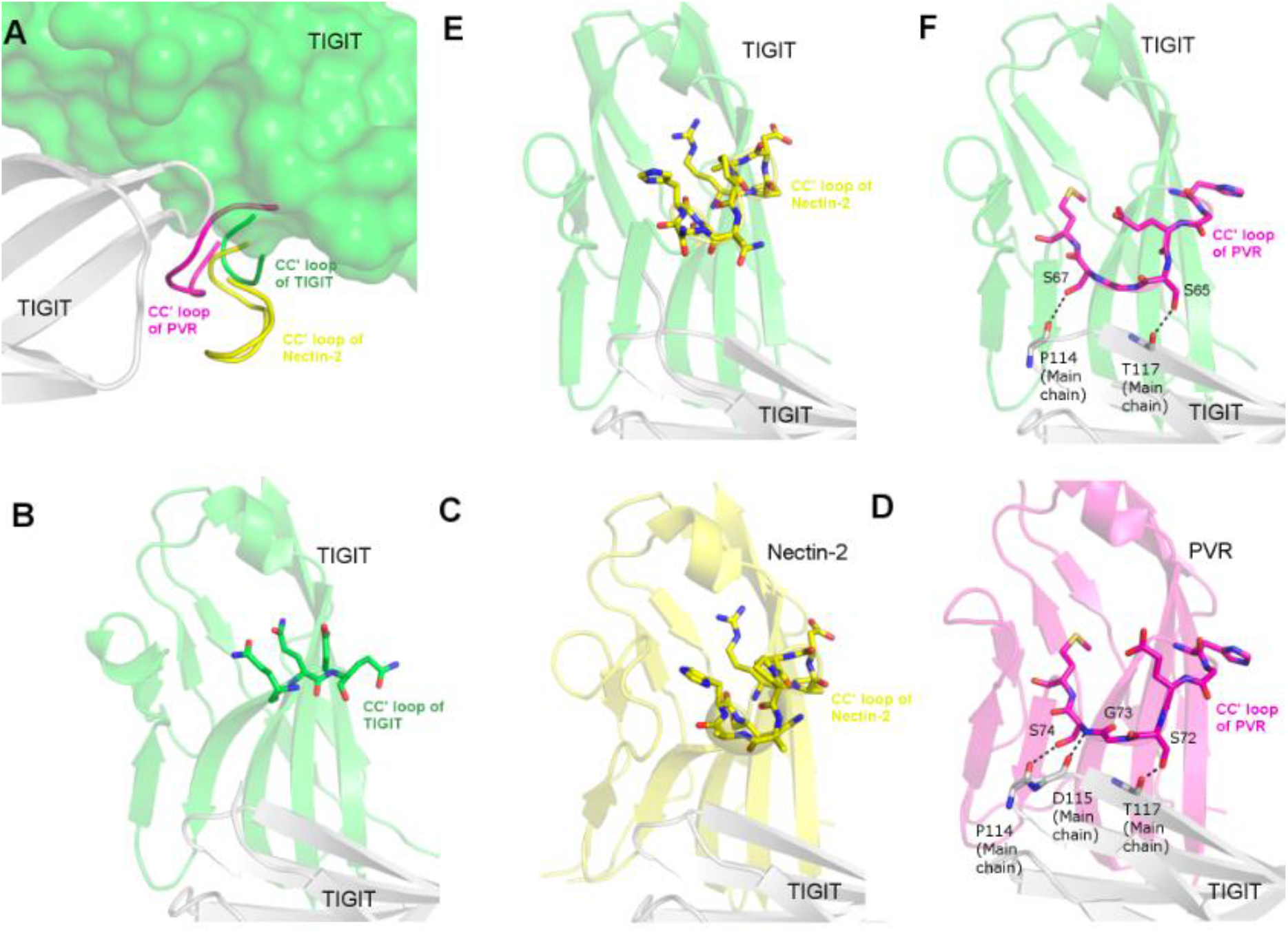
Modification of CC′ loop might lead to increase in binding affinity: Panel A illustrates the difference in the CC′ loops of TIGIT (ribbon, green), Nectin-2 (ribbon, yellow) and PVR (ribbon, magenta) when bound to TIGIT (cartoon, light grey). Since, all the three ligands have the same mode of interaction, TIGIT:Nectin-2 and TIGIT:PVR complexes were superposed on TIGIT(surface, green):TIGIT homodimer. Panel B, C and D illustrates the mode of interaction from experimentally determined structures of TIGIT (cartoon, green):TIGIT (cartoon, light grey), Nectin-2 (cartoon, yellow):TIGIT (cartoon, light grey) and PVR (cartoon, magenta):TIGIT (cartoon, light grey). The CC′ loops of the respective structures are shown in sticks. H-bond interactions of CC′ loop of PVR with TIGIT is indicated with black dotted line. Panel E and F depicts the best docked poses of T_N_:WT-T and T_P_:WT-T respectively where, modified TIGIT in both E and F panel in represented in green, cartoon and the spliced CC′ loops of Nectin-2 and PVR are represented in sticks, yellow and magenta respectively. Wild type TIGIT in both panels E and F is represented in light grey, cartoon. The interacting residues of T_P_ and TIGIT in panel F are represented in sticks.

Experimentally determined structures of TIGIT:PVR (PDB:3UDW) and TIGIT:Nectin-2 (PDB:5V52) and TIGIT:TIGIT homodimer (PDB:3RQ3) were superposed in the crystallographic modelling program *Coot*. In two independent attempts, CC′ loops of one of the TIGIT monomer of the TIGIT homodimer (PDB:3RQ3) was swapped with the CC′ loop of Nectin-2 and PVR. The complex structure thus created comprising of TIGIT with the CC′ loop of Nectin-2 (named T_N_) and wild type TIGIT (WT-T) was subjected to energy minimization using UCSF Chimera with Amber ff14SB forcefield. The same was done with the heterodimer containing TIGIT with the CC′ loop of PVR (T_P_) and WT-T. Since, the mode of binding of TIGIT:TIGIT homodimer and TIGIT:PVR/Nectin-2 heterodimers are similar, the resulting energy minimized structures of WT-T:WT-T, T_P_:WT-T and T_N_:WT-T were docked using RosettaDock via ROSIE server.

To examine the veracity of the docking algorithm, docking trials were also performed on experimentally determined TIGIT:TIGIT, TIGIT:Nectin-2 and TIGIT:PVR complexes. It was observed that the interface score was a more accurate parameter than the docking score, when the scores were compared with the docked poses which were in accordance (RMSD<1) with the experimentally determined structures. Hence, these interface scores (more negative score, better the strength of interaction) were used as a reference in order to compare with that of the modified heterodimers (T_P_:WT-T and T_N_:WT-T). Results showed that the WT-T:WT-T had an interface score of −8.2, whereas WT-T:Nectin-2 and WT-T:PVR had scores of −9.6 and −9.3 respectively. The best docked poses of T_N_:WT-T and T_P_:WT-T gave interface scores of −8.4 and −9.1 respectively (Figure 6E and 6F respectively). It is interesting to note that difference in the interface scores of T_N_:WT-T and WT-T:WT-T (−0.2) is higher than that between T_P_:WT-T and WT-T:WT-T (−0.9), which indicates that the contribution of the CC′ loop of PVR plays an important role in the interaction interface than the CC′ loops of TIGIT and Nectin-2. It can be expected that the differences in interaction due to other front face residues of Nectin-2 and PVR also contribute to the observed differences in affinity between TIGIT:Nectin-2 and TIGIT:PVR complexes. However, the docked T_N_:WT-T and T_P_:WT-T structures are similar to the experimentally determined (PDB:3UDW and 5V52 respectively) structures and maintain the similar interaction seen between the CC′ loops of PVR and Nectin-2 with the WT-T. The interface scores are reflective of the fact that CC′ loop of Nectin-2 makes very few hydrophobic interactions, whereas CC′ loop of PVR makes significant amount of hydrophobic and hydrophilic interactions, which is also observed in the experimentally determined structures. This increase in the interaction score of T_P_:WT-T is due to the additional interactions made by the CC′ loop of PVR with the cognate TIGIT monomer, similar to what is seen in TIGIT:PVR. Conversely, it was observed that when the CC′ loop of PVR was replaced by the CC′ loop of TIGIT (P_T_) and docked with the WT-T in the canonical orientation, there were loss of important interactions which is also supported by low interface score of −6.4. These interface scores indicate that the modifed versions of TIGIT (T_P_ and T_N_) have a higher binding affinity with its corresponding monomer of TIGIT (WT-T) than wild-type TIGIT itself. Interestingly, mere increase in the length of the CC′ loop doesn’t translate into increased binding affinity as the length of CC′ loop of PVR is shorter than that of Nectin-2. Functionalities of the residues present in the CC′ loop and their interactions with the cognate ligand appear to take precedence. Commensurate with this observation, experimental swapping of CC′ loop of Nectin-2 with the CC′ loop of PVR, resulted in a Nectin-2 mutant that showed higher affinity towards TIGIT than wild type Nectin-2 (Deuss et al., 2017). These results might provide the motivation to proceed with wet lab experiments to confirm our hypothesis with not only TIGIT/Nectin pathway, but other medically relevant ICRs as well. It must also be noted that the residues of the CC′ loops of TIGIT, Nectin-2 and PVR are not highly conserved (data not shown). Hence, these modifications to improve the binding affinity could be done with limited fear of structural instability or non-specific binding to undesired targets.

In conclusion, the immune checkpoint blockade has brought about a paradigm shift in cancer therapy, however, rising reports of toxicity of mAbs coupled with its high cost, suspected resistance, requirement of high dose, poor solid tumour penetration and so on, have pushed for alternatives that can mimic the function of mAbs. Since, the soluble isoforms of these ICRs are naturally used by our immune system to tweak the immune response, rationally designed naturally occurring ICRs to mimic the function of therapeutic mAbs appears to be an effective approach. However, to design such a protein, sound structural information of the proteins and intuitive modification strategies are critical. In lieu of the above concept, the CC′ loop of the IgV containing ICRs have been identified as the prime target for modifications in this study. The CC′ loop appears to influence the binding affinity of IgV domain containing ICRs in about half the ICR complexes analysed. In about quarter of these complexes, the CC′ loop contributes more towards the interaction-interface than the FG loop. The CC′ loop is flexible enough to accommodate the incoming ligand and once bound the CC′ loop acts like a clamp to increase the binding affinity of the complex. In a few cases, the CC′ loop takes a preformed non-canonical conformation in order to bind to small molecules as elucidated in the case of mTIM-3 and CD22. These observations highlight the versatility of the CC′ loop in order to form a stable complex and provide additional affinity to the interaction. Since, most of the residues involved in CC′ loop are not highly conserved (data not shown), educated modifications on the CC′ loop to increase the binding affinity, might raise limited concerns on protein instability and cross-reactivity. This hypothesis was put to the test with *in silico* modifications of TIGIT, where the CC′ loop of TIGIT was replaced with CC′ loops of Nectin-2 and PVR. The results indicated trends which suggests that modified TIGIT (T_N_ and T_P_) bound to wild type TIGIT with higher affinity, when compared to the canonical TIGIT homodimer. This work attempts to provide novel insights for rational design of soluble ICRs to mimic the role essayed by mAbs with an expected low toxicity and cost.

## Methodology

### Retrieving structures and complexes for the study

ICR complexes and other immune related receptors which contained at least one IgV domain were mined from literature (Pardoll, 2012) and their apo structures and complexes were retrieved from worldwide protein databank (wwPDB) (Berman et al., 2003). The presence of IgV domain in these retrieved structures were confirmed by InterPro-Protein sequence analysis and classification (Finn et al., 2017). The secondary structure description as provided by PDB header was considered while determining the identity and length of the CC′ loop. The residue numbering in the PDB coordinate file was considered for the analysis and the same was used in the manuscript.

### Structure and interaction analysis

The interactions amongst the ICRs and their ligands were analysed using PDBe PISA (Krissinel and Henrick, 2007), where H-bond length not greater than 3.5Å and not smaller than 2.5Å were considered. Van der Waals interactions were considered if the two residues were separated by a distance of less than 5Å. Coot (Emsley et al., 2010) and PyMol (Delano, 2004) were used to visualize the complexes and their interactions. Buried surface area (BSA) of each monomer in the complex was considered for calculating the contribution of CC′ loop of the respective monomer which is indicative of Van der Waals interactions. The post-interaction movement of CC′ loop was considered significant, if the maximum deviation of the C_α_ atom of any residue which belongs to CC′ loop is more than 3Å. The sequence logos highlighting the conservation of the residues of CC′ loop have been performed using WebLogo (Crooks et al., 2004).

### *In silico* modelling and energy minimization trials

*In silico* splicing of the CC′ loops of PVR and Nectin-2 onto one TIGIT monomer of the TIGIT homodimer was performed using Coot using experimentally solved structure of TIGIT homodimer in hexagonal space group (PDB: 3RQ3) as the template structure. The CC′ loops of PVR and Nectin-2 were excised and replaced the CC′ loop of TIGIT by superposing TIGIT homodimer structure with TIGIT:PVR (PDB:3UDW) and TIGIT:Nectin-2 (PDB:5V52) heterodimers independently. These *in silico* modified heterodimers of TIGIT (wild type TIGIT with TIGIT with CC′ loop of Nectin-2 or PVR) were subjected to energy minimization using UCSF Chimera’s Amber ff14SB forcefield (Pettersen et al., 2004). These values were documented for further analysis.

### Protein-protein docking trials

As mentioned earlier, the mode of interaction of TIGIT homodimer and heterodimers are the same. Hence, the modified TIGIT heterodimers were docked using RosettaDock (Chaudhury et al., 2011; Lyskov and Gray, 2008) via ROSIE server (Lyskov et al., 2013). Since, the modified heterodimers were derived from the exact pose of the experimentally determined structure (PDB:3RQ3), interface score was given paramount importance. The interface score was found to be more relevant when the docked poses were compared with experimentally determined structures. This parameter would help in inferring the trend associated with the impact of modification of the CC′ loop. The docked pose with the highest interface score which had a RMSD<1 with the submitted template structure, was considered for further investigations. Negative interface score indicates stronger interactions between the interfaces of the two proteins. Furthermore, thorough structural analysis was carried out on the selected docked pose to ascertain the number and type of interactions made by the CC′ loop with its binding partner. These interactions were compared with the interactions made by the CC′ loops of PVR and Nectin-2 with TIGIT as seen in experimentally determined structures (PDB:3UDW and 5V52 respectively) using PDBe PISA (Krissinel and Henrick, 2007) and PIC (Tina et al., 2007).

## Abbreviations

ICRs: Immune checkpoint receptors
IgV: Immunoglobulin V domain
mAbs: monoclonal antibodies

## Acknowledgment

UAR would like to thank DBT Ramalingaswamy fellowship for his fellowship.

## Author contribution

UAR conceptualized, supervised and reviewed the manuscript. SVK performed the analysis and UAR and SVK wrote the manuscript

